# Anatomical connectivity profile development constrains medial-lateral topography in the dorsal prefrontal cortex

**DOI:** 10.1101/2022.02.07.479322

**Authors:** Wen Li, Weiyang Shi, Haiyan Wang, Jin Li, Yue Cui, Kaixin Li, Luqi Cheng, Yuheng Lu, Liang Ma, Congying Chu, Ming Song, Zhengyi Yang, Tobias Banaschewski, Arun L.W. Bokde, Sylvane Desrivières, Herta Flor, Antoine Grigis, Hugh Garavan, Penny Gowland, Henrik Walter, Rüdiger Brühl, Jean-Luc Martinot, Marie-Laure Paillère Martinot, Eric Artiges, Frauke Nees, Dimitri Papadopoulos Orfanos, Herve Lemaitre, Tomáš Paus, Luise Poustka, Sarah Hohmann, Sabina Millenet, Juliane H. Fröhner, Lauren Robinson, Michael N. Smolka, Jeanne Winterer, Robert Whelan, Gunter Schumann, Lingzhong Fan, Tianzi Jiang, IMAGEN Consortium

## Abstract

The prefrontal cortex (PFC) is a highly variable, evolutionarily expanded brain region that is engaged in multiple cognitive processes. The subregions of the PFC mature relatively late compared with other brain regions, and the maturation times vary between these subregions. Among these, the dorsomedial and dorsolateral prefrontal cortex (dmPFC and dlPFC) share a parallel topographic pattern of functional connectivity, while participating in different types of complex behaviors. However, the developmental trajectories of the two areas remain obscure. In this study, we uncovered differences in the developmental trends of the dmPFC and dlPFC. These differences were mainly caused by structural and functional changes in the medial area of the superior frontal gyrus (SFG). The developmentally different arealization patterns were verified using multiple parcellation approaches with multimodal data, including structural magnetic resonance imaging (sMRI), diffusion MRI (dMRI), resting state functional MRI (rfMRI), and a publicly available transcriptomic dataset. Human brain gene expression data was also used to perform downstream analyses, which could inform us about the potential biological mechanisms underlying the developmentally different arealizations. Furthermore, behavioral analyses hinted at the effects of regionalization on ontogeny. In brief, this study revealed a tendency toward a medial-lateral prefrontal division and can provide a fuller understanding of the potential underlying genetic underpinnings as well as of the potential effects on developmental behavior.

## Introduction

The dorsomedial and dorsolateral prefrontal cortex (dmPFC and dlPFC) are central hubs of cognitive control and decision-making ^1,2^. The dmPFC is implicated in response conflict detection ^3,4^ whereas the dlPFC is more involved in adjusting behavior ^5^. On the other hand, the dmPFC and dlPFC are also recognized as having parallel functional topographies, which are manifested as a posterior-to-anterior connectivity gradient ^5,6^. Processing endogenously and exogenously relevant tasks also involves distinct anterior prefrontal networks along a medial lateral axis ^7^. This functional segregation of medial and lateral areas is well demonstrated in neuroimaging studies, mostly in adult cohorts. However, the prefrontal lobe is one of the last regions of the brain to mature ^8^ in that the processes of synaptic pruning, gray matter decrease, and white matter augmentation continue into adulthood ^9,10^. Thus, when the functional separation of the dmPFC from the dlPFC begins and which factors drive this separation during development remain poorly understood.

Identifying the connectivity patterns in various developmental stages has broad implications for our fundamental understanding of the formation of the topological organization patterns of brain regions ^11^. Because the general consensus is that the cortical subregions can be defined as clusters of voxels that share similar connectivity patterns, anatomical connectivity profiles derived from diffusion MRI (dMRI) have been used to parcellate the human brain ^12,13^. In addition, the functional diversity of brain regions is constrained by structural connections ^14^, and structural connectivity patterns can also predict functional activation ^15,16^. Moreover, broad behavioral changes during development are accompanied by major changes in anatomical and functional brain networks ^17^, and some researchers found that the segregation of brain connections shaped and constrained the segregation of brain functions across the lifespan. For instance, Qin and his colleges demonstrated that the less segregated functional network of the subregions of the amygdala in children led to weaker affective functions compared with adults ^18^. Age-related variation in connectivity patterns during memory formation was also reported ^19^. Hence, our goal was to reveal the functional separation of the dmPFC from the dlPFC in developmental stages from childhood to early adulthood using anatomical connectivity contrasts for the purpose of increasing our insight into the differences in connectivity patterns and the extent of separation between different subregions.

However, which factors drive the separation of connections and functions between subregions warrants more genetics-related work. It is well recognized that the spatiotemporal regulation of genetics plays an important role in the arealization of distinct cortical areas ^20^. These distinct areas are influenced by genes that exhibit highly regionalized expression patterns ^21^; thus, the cortical areas could be parceled into genetic subdivisions with meaningful structural and functional region boundaries using only genetically informative data at either a coarse level ^22^ or a finer scale ^23^. In addition, the connectivity patterns are mediated by the gene expression pattern. Zhao et al. reported that the white matter microstructure showed that it is highly heritable ^24^, and the regulatory effect of gene expression on the formation of functional connections has also been reported ^25^. Furthermore, the substantial maturation of cognitive control processes during adolescence are affected by genetic factors ^26^. Exploring these processes can allow for a better understanding of the changes that occur during the arealization of the brain cortex as well as during the maturation process associated with cognition and behavior.

In this work, we developed an innovative developmental atlas of the superior frontal lobe from adolescence to early adulthood using Lifespan Human Connectome Project in Development (HCP-D) data and employing a systematic, validated parcellation pipeline. To our knowledge, this is the first detailed version of a developmental atlas that depicts the changes in the arealization pattern of the frontal lobe during human development and can, therefore, provide a reference and resource for neurodevelopmental studies. We validated our findings using the IMAGEN dataset, which is a representative longitudinal dataset and uses information extracted from multimodal neuroimaging. Both structural and functional connectivity patterns were compared separately for different age brackets. In addition, the gene expression maps provided by the Allen Human Brain Atlas (AHBA) were used to explore the underlying fundamentals that modify this segregation. By linking gene expression to age-related structural and functional connections, we provided evidence for a possible mechanistic substrate by which gene expression could regulate the regionalization patterns by regulating a battery of physiological processes. In addition, we explored the relationship between the behavior and the similarity of the functional connections between subregions.

## Methods

The data used in this article were from HCP-D and IMAGEN. After preprocessing, the dMRI data were used for the parcellation, and multimodal data were used to verify the appropriate patterns. Gene expression and ANOVA analyses were performed to further explore whether subregion division is associated with age and to discover its biological fundamentals. The relationship between behavior and connections was explored using PLS regression. Finally, how the functional connections affect behavior was also investigated. The detailed description is as follows.

### Participants

The primary dataset used in this work was the HCP-D ^27^. The IMAGEN dataset was then used to verify the credibility of our results and, because it is a longitudinal dataset, it was also used for the behavioral analysis. The basic information about the two datasets is described below.

#### HCP-D data

The HCP-D Release 1.0 includes T1 weighted (T1w) MRI, resting state functional MRI (rfMRI), task fMRI, and diffusion MRI (dMRI) data for 655 subjects (323 males, age 5-21) acquired at four sites across the USA. Two separate MRI sessions were acquired on a 3-T Siemens Prisma platform using a 32-channel head coil. A battery of cognitive tasks and self-reports were also performed. Multi-echo MPRAGE ^28^ was used to acquire the T1w scans at a resolution of 0.8 mm. The dMRI data consisted of 185 directions on 2 shells of b = 1500 and 3000 s/mm^2^ along with 28 b = 0 s/mm^2^ images in four consecutive dMRI runs at a resolution of 1.5 mm. The rfMRI scans used a 2D multiband (MB) gradient-recalled echo (GRE) echo-planar imaging (EPI) sequence (MB8, TR/TE = 800/37 ms, flip angle = 52°) ^29^, and the resolution was 2.0 mm.

After consulting the literature from international well-known child research organizations and related research across the life span, we split the data into four age groups from child to adult and termed them post-childhood (8-10y), early-adolescence (11-14y), post-adolescence (15-17y), and young-adulthood (18-21y) (Table S1). Subsequent group analyses were based on this division. All of the partitioning methods applied in this study were the same for all age brackets.

#### IMAGEN data

The multi-site longitudinal IMAGEN project involved more than 2000 adolescents ^30^. The participants participated in the first data collection at about 14 years old with a follow-up imaging collection in their 19^th^ year. Brain imaging data were collected at eight sites on 3T MRI systems made by four manufacturers (Siemens, Philips, General Electric, and Bruker). The scanning parameters and sequence protocol were devised for each image-acquisition technique and held constant across sites. To ensure the comparability of the fMRI data from the different sites, the deviation across sites was controlled by adding it as a nuisance covariate in our analyses. The T1w scans were acquired with a gradient-echo MPRAGE sequence (1.1 mm slice thickness, TR = 2,300 m, TE = 2.8 m). The diffusion tensor images were acquired using an Echo Planar imaging sequence (4 b-value = 0 s/mm^2^ and 32 diffusion encoding directions with b-value = 1300 s/mm^2^; 60 oblique-axial slices (angulated parallel to the anterior commissure/ posterior commissure line); echo time ≈ 104 ms; 128 × 128 matrix; field of view 307 × 307mm; voxel size 2.4 × 2.4 × 2.4 mm), adapted to tensor measurements and tractography analysis. Where available, a peripherally gated sequence was used; when this was not possible, TR was set to 15s, approximately matching the effective TR of the gated scans. The functional runs used a gradient-echo echo-planar imaging (EPI) sequence with echo time = 30 ms, repetition time = 2200 ms (2.4 mm slice thickness), flip angle = 750, in-plane resolution = 3.4 × 3.4 mm, and field of view = 220 × 220 mm.

### Preprocessing of HCP-D and IMAGEN

The pre-processing pipeline of HCP-D used in our study (v4.1.3), was similar to that described in the minimal preprocessing pipelines for HCP Young Adult (HCP-YA) ^31^, but a few adjustments were made to adapt to the children’s data (see https://github.com/Washington-University/Pipelines). The way that the multimodal data was pre-processed is as follows.

The preprocessing procedures for the structural data consists of the PreFreeSurfer-pipeline, the FreeSurfer-pipeline, and the PostFreeSurfer-pipeline. The PreFreeSurfer-pipeline was used to create native volume space by averaging the T1w and T2w images separately, correcting the bias field, performing brain extraction, and aligning the T1w or T2w images to a specified MNI template. The FreeSurfer-pipeline that we used was approximately that described by Fischl 32 but employed a slightly improved algorithm to adapt the scan protocol. The PostFreeSurfer-pipeline converted the data to the NIFTI and GIFTI formats and generated a final brain mask and myelin maps in addition to Connectome Workbench spec files in surface space.

The diffusion-pipeline began with normalizing the b0 intensity followed by estimating the EPI distortion, eddy current distortions, and subject head motion into a Gaussian process predictor to allow for correction. Then the b0 image was registered to the T1w image using boundary-based registration. Finally, diffusion data were registered to native structural space and masked to the appropriate size. An additional step, FSL’s BEDPOSTX, was applied to estimate the uncertainty of each fiber in each voxel for three possible fiber orientations ^33^.

The pre-processing of the rfMRI data included the fMRI-Volume-pipeline and the fMRI-Surface-pipeline. The former consisted of motion correction, EPI distortion correction, and concatenating all transformations into a single nonlinear transformation. The latter mapped the volume time series to the standard CIFTI gray ordinates space by methods including, but not limited to calculating the partial volume weighted ribbon-constrained mapping algorithm, excluding voxels that had a locally high coefficient of variation in the time series data, and utilizing the geodesic Gaussian surface smoothing algorithm (with full-width half-maximum (FWHM) of 2 mm). Finally, the rfMRI data were further processed using ICA-FIX ^34^ to remove structured artifacts.

For the IMAGEN dataset, high-resolution anatomical MR images were warped to common space using linear and nonlinear transformations provided by SPM12. The FMRIB Diffusion Toolbox in FSL (www.fmrib.ox.ac.uk/fsl) was applied to preprocess the diffusion data. To reduce the impact of the head motion and eddy currents on the subsequent analysis, affine registration to the first b=0 image was executed. Then the FSL brain extraction tool (BET) was used to extract the brain ^35^. BEDPOSTX was used as described above.

Because of a lack of sufficient rfMRI data, the face-related fMRI data were modified to emulate the resting state data. The event-related signals were regressed from the face-related fMRI data, and the emulated resting state data were derived from the residuals ^36^. Then, after slice-timing correction and non-linear registration into MNI space, the rfMRI data were resampled to 3 × 3 × 3 mm resolution. To control the confounding effects of motion artifacts on the rfMRI data, we used head motion parameters (3 rotations and 3 translations). Additional preprocessing included detrending, bandpass filtering (0.01−0.1 Hz), global signal regression, white matter regression, regressing the mean signal from the cerebrospinal fluid, and smoothing (FWHM = 5 mm) on the surface. Scans in which the framewise displacement (FD) was > 0.5 mm were scrubbed ^37^. The data of the subjects who had fewer than half of their volumes remaining after the scrubbing or those with excessive head motion (defined a priori as >3 mm translation or > 3° rotation) were excluded in this study ^38^.

### Connectivity-based parcellation

Creating a group template and defining the initial seed mask are the first steps toward establishing a credible parcellation. We adopted the asymmetric NIHPD pre-to-early puberty (7-11y), early-to-advanced puberty (10-14y), post-puberty (13-18y), and MNI152 adult templates ^39,40^ which correspond to the ages of 8-10y, 11-14y, 15-17y, and 18-21y, respectively. The skull-stripped T1 image for each subject belonging to each age group was registered to the corresponding templates by nonlinear transformations and then merged to generate a group template of every age bracket after averaging. The initial seed masks came from Fan, et al. ^13^ and were used as regions of interest (ROIs) for the subsequent parcellation.

Then, 5000 streamline fibers for each voxel in the SFG were sampled to estimate the connectivity probability using the FSL package ^33^. Next, the results of fiber tracking were extracted and down-sampled to 5 mm isotropic resolutions to constitute a connectivity matrix ^41^, in which the rows each represent one voxel in the SFG and the columns are the voxels of the whole brain. Then, the connectivity matrix was transformed into symmetric cross-correlation M-by-M matrixes, in which M represents the numbers of voxels, which are treated as features for the subsequent clustering. Finally, the connectivity matrix for each subject was entered into the automatic parcellation pipeline, and maximum probability maps were created across all the subjects in each age bracket ^13,42^. Various clustering methods, including spectral clustering, k-means clustering, and SIMLR clustering, were tested to divide each ROI into 2-10 subregions, and similar parcellation patterns were obtained. The parcellation results presented in this article were from the spectral clustering algorithm.

Determining which areas correspond to homologous areas in the different age groups was a very difficult aspect of this developmental research. A potential solution for matching regions was to measure the degree of overlap between regions after registering them to the same template. The higher the degree of overlap the greater the possibility that the regions were homologous. However, it was obviously not accurate to determine the homologous region only by overlap degree because the connection patterns also need to be taken into account 14. In this study, therefore, we also took the similarity of the connections into consideration. Due to a lack of whole brain atlases for all the age brackets, we used the 72 major white matter tracts specific for each individual as the target areas. These tracts were acquired by segmenting white matter tracts in fields of fiber orientation distribution function peaks using a fully convolutional neural network ^43^. The advantage to adopting both overlap degree and connection similarity to match regions is to reduce the errors caused by registration and parcellation between different groups.

### Determining the optimal number of clusters

There is no gold-standard for determining the degree of validity of a given parcellation method of brain images because researchers lack biological ground truth on which to base their measures of validity ^11^. For this reason, we used as many kinds of assessment methods as possible to determine the optimal number of clusters for each region. Since the parcellation pattern for a group should be adaptable to a large number of individuals, we employed the Dice index to evaluate the parcellation consistency among individuals. The Dice coefficient describes the degree of overlap between different individuals of the same region. Counting the ratio of the variances between and within clusters is also important for deciding whether a clustering method is successful in that a good clustering method should have large inter-group differences and small intra-group differences. NMI ^44,45^ and Cramer’s V (CV) 46 were both used to measure the topology consistency. NMI quantifies the amount of information received from one partition to another, while CV characterizes the similarity between the two partitioning modes. Another index, the silhouette index (SI) ^47^, was used to describe the similarity of a voxel to its own cluster compared with other clusters. We also assessed the individual consistency (IC) of the connectivity patterns for different cluster numbers. The connectivity patterns were defined as the 1×246 vectors in which every value represents the normalized sum of the number of fiber bundles from the cluster to the target areas after removing the false-positive connections ^42^. The target areas were defined by the Brainnetome Atlas, in which the whole brain was parceled into 246 subregions ^13^. To display the IC results more intuitively, we regressed the decreasing trend resulting from the increase with age.

### Multimodal validation

Data from other modalities were also used to validate the parcellation. For the parcellation made using the fMRI data, another popular parcellation method, boundary mapping, was used to verify and supplement the parcellation. Unlike the cluster method, the boundary mapping approach used a local border detection technique to detect a border by locating the most abrupt spatial changes in the assessed feature ^11^. Sharp changes in gradient values were considered to be indicative of potential boundaries. The boundary mapping used resting-state functional connections, which were defined as the Pearson correlation between voxels from the seed area and the whole brain, to define the parcels. The partition algorithm used in this article was watershed by flooding. This method defined regions in the gradient maps by starting from the local minimum and grew iteratively until it reached a position that could be vaguely assigned to multiple regions ^48^. With respect to cortical thickness, the feature values did not vary sharply enough across regions to be used to define clear boundaries; therefore, gradients of thickness were calculated to observe how the boundaries of the subregions changed with age ^49^.

To explore the detailed morphology and architecture of the areas of interest, BigBrain ^50^ was utilized. Based on the parcellation results, the two regions with the most pronounced changes in parcellation patterns during development, the A9l and A9m areas, were chosen for the subsequent architecture study. In both areas, the distribution of all the values, which indicated the density of cells and included information about the types of cells, was found to be able to distinguish the two areas.

### ANOVA analyses between age brackets

ANOVA analyses were also used to examine the differences between the connections of the two areas quantitatively. For each subject in each age bracket, we computed the structural connection of each subregion as a proportion of the connections with all the subregions of the ROI, to explore the transition in the connections for each subregion during development. Then the proportion was compared between the age groups using two-way ANOVAs separately for each subregion with ‘age’ and ‘gender’ as fixed and random factors, respectively. For both the white matter connections and the functional connections, the 246 subregions defined by the Brainnetome Atlas were applied as target areas. Additional target areas, the 7 networks areas from Yeo et al. ^51^, were adopted to further explore the transition in functional connectivity development. Then, a paired *t*-test analysis was performed between the groups to further explore the changes in connection patterns. Both the ANOVA and paired *t*-test analyses were performed using Matlab.

### Differential gene analysis and gene enrichment analysis

Regional microarray expression data were obtained from 6 post-mortem brains (1 female, ages 24.0-57.0) provided by the Allen Human Brain Atlas (AHBA, https://human.brain-map.org). The data were processed using the abagen toolbox (version 0.1.1; https://github.com/rmarkello/abagen). First, the microarray probes were reannotated using data provided by ^52^; the probes that did not match a valid Entrez ID were discarded. Next, the probes were filtered based on their expression intensity relative to background noise ^53^, such that probes with intensities less than the background in ≥50.00% of the samples across the donors were discarded, yielding 31,569 probes. When multiple probes indexed the expression of the same gene, we selected and used the probe with the most consistent pattern of regional variation across the donors (i.e., differential stability ^54^). In this context, the regions corresponded to the structural designations provided in the AHBA ontology. The MNI coordinates of the tissue samples were updated to those generated via non-linear registration using Advanced Normalization Tools (ANTs). The samples were assigned to brain regions in the provided atlas only if their MNI coordinates were directly within a voxel belonging to a parcel. To reduce the potential for misassignment, sample-to-region matching was constrained by hemispheric and gross structural divisions (e.g., a sample in the left cortex could only be assigned to an atlas parcel in the left cortex ^52^).

All tissue samples not assigned to a brain region in the provided atlas were discarded. Inter-subject variation was addressed by normalizing the tissue sample expression values across the genes using a robust sigmoid function from Fulcher, et al. ^55^. Normalized expression values *xnorm* were then rescaled to the unit interval: *x_scaled_ = x_norm_ – min(x_norm_))/(max(x_norm_)-min(x_norm_))*. Gene expression values were then normalized across tissue samples using an identical procedure. Normalization was performed separately for samples in distinct structural classes (i.e., cortex, subcortex, and cerebellum). Samples assigned to the same brain region were averaged separately for each donor and then across the donors, yielding a regional expression matrix with 235 rows, corresponding to brain regions (11 ROIs were discarded because of having no samples, see Table S2), and 15,605 columns, corresponding to the retained genes.

The differential gene analysis of the A9l and A9m areas was performed using JuGEx, which integrates the frameworks of the AHBN and Julich-Brain atlases. The processed expression data were extracted for the two subregions. Next, a single-probe analytical mode was chosen to investigate the strongly expressed genes. N-way ANOVA permutations and multiple comparisons with a family-wise error (FWE) correction were included in the statistical analyses. Finally, the progression of the dynamic gene expression of the genes that changed throughout childhood was explored from infancy to old age, supplemented by data from the Human Brain Transcriptome (HBT, http://hbatlas.org/) ^56^.

Gene Set Enrichment Analysis (GSEA) can be used to discover which specific biologically significant gene sets are significantly related to the expression or other measurement levels of samples from selected areas. Because it is not limited to differential genes, it is easier to discover the impact of subtle gene set differences on biological processes ^57,58^. The gene samples from the A9m and A9l areas together with their gene expression vectors were extracted based on the AllenBrain atlas coordinates. Because we lacked strong prior knowledge about gene enrichment in the two areas, all of the gene sets provided by GSEA were employed to detect the enrichment of genes. Then, the significant enriched gene sets were visualized using Cytoscape ^59^.

### Coupling between connections and gene expression

To obtain the age-related-connection matrix, the dependence of the changes in both structural and functional connections on age were calculated using the Spearman correlation. Next, PLS regression was used to explore the relationship between the age-related-connection matrix and the gene regional expression matrix. In the PLS regressions, gene expression data were used as predictor variables to assess the changes in the age-related connection matrices separately for structural and functional connectivity. Each component of the PLS was the linear combination of the gene expression values. This process was followed by a validity check. The null hypothesis, that the PLS1-explained covariance between the connection matrix and the whole-genome expression is not larger than the expected accidental covariance, was then tested using permutation tests (10,000 iterations).

Bootstrapping was used to estimate the variability of each gene’s valid component and to rank the genes according to their contribution to the PLS. Both positive and negative genes with an FDR less than 5% were listed and followed by a gene enrichment analysis using GORILLA ^60^. The gene ontology (GO) terms they were related to were projected to two-dimensional space and visualized as semantic similarity-based scatterplots by REVIGO ^61,62^.

### Relationship to behavior

To explore the behavioral differences between the two subregions, A9l and A9m, NeuroSynth was used for meta-analytical decoding ^63^. This provided a platform for automatically synthesizing the results from 14371 neuroimaging studies. Thousands of behavioral or psychological terms were abstracted according to their frequency in these articles. The first ten terms most relevant to each subregion were extracted to reveal the differences between A9l and A9m in the behavioral domain.

The functional connection matrix, which is used for behavior-related study, was defined as the Pearson correlation coefficients of pairs of nodes, where the nodes were parcels identified in the Brainnetome atlas. In each node, the time series of each voxel was normalized to *z* scores using the mean and standard deviation for each subject; then the voxels within the same node were averaged. The similarity of the connectivity patterns of the A9m and A9l from the same 14y individual was then computed.

In the next step, the potential dependency between the similarity and behavior were studied based on IMAGEN data. Only subjects with complete demographic information, including age, gender, handedness, acquisition site, and related behavioral data were included in the behavioral study. The SFG is known to be associated with delayed reward systems ^64,65^, and the most widely used scale to measure this behavior is the Monetary Choice Questionnaire (MCQ), which is a self-management questionnaire consisting of 27 items. The participants were required to choose between a smaller, immediate monetary reward and a larger, delayed monetary reward in each item. Then the scores were fitted using a hyperbolic function model, Kind(*K*) = ((LDR/SIR)-1)/DELAY, where SIR refers to short immediate rewards, LDR represents long delay rewards, and DELAY is the number of days of delay. Smaller values indicate a lack of discounting and a preference for delayed rewards, and higher values indicate strong discounting and a preference for immediate rewards. Thus, higher values of *K* are indicative of high levels of impulsivity ^30,66,67^. The partialcorr function in Matlab was used to model the relationship between the functional connections similarities and the behavior changes, which were defined as the behavioral difference values between adolescences and adults, while regressing the concomitant variables of age, handedness, site, and economic condition of each subject. Permutation tests (10,000 iterations) were then performed for the statistical analysis.

**Figure 1.**
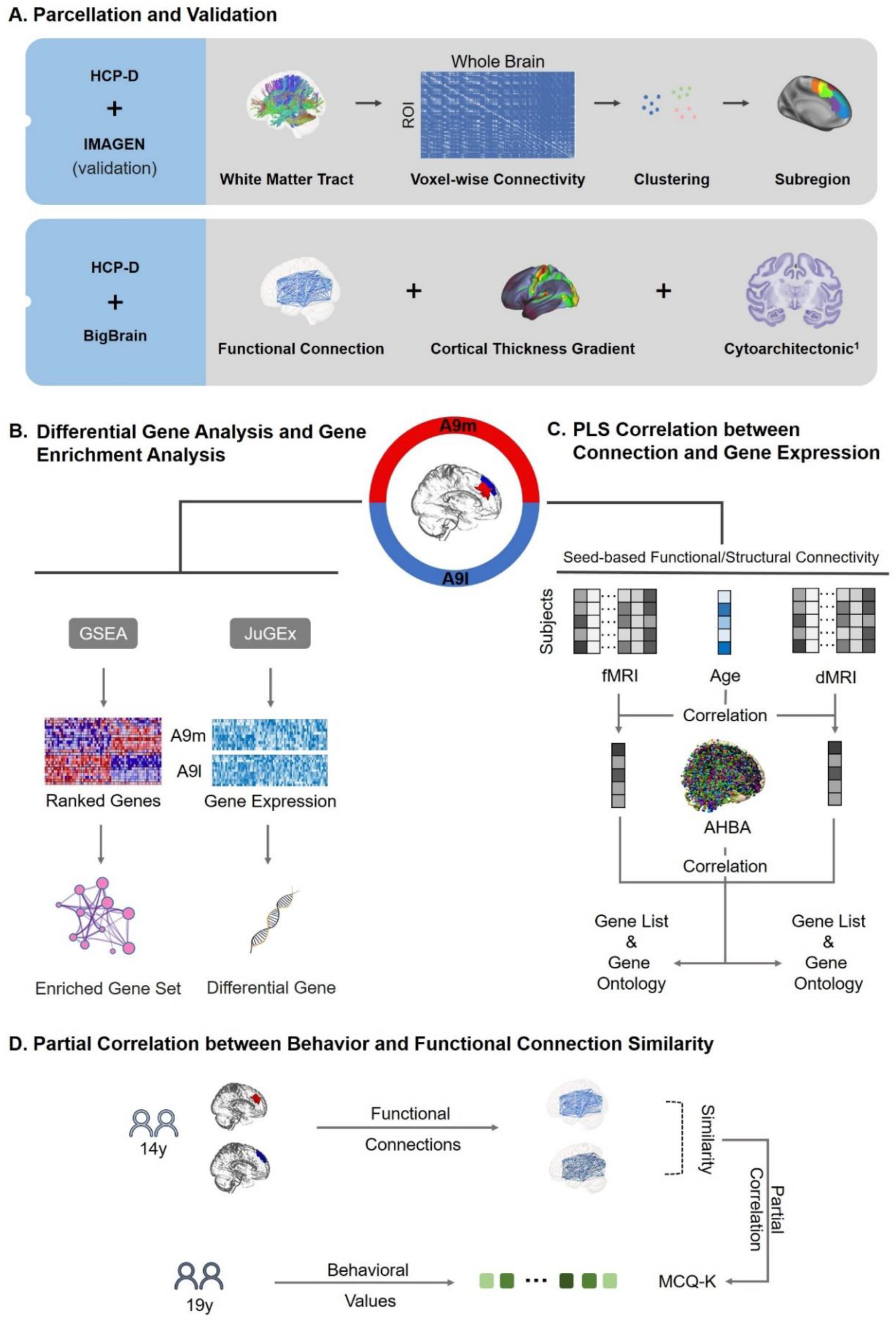
Summary of the analysis flowchart. **(A)** Framework of the parcellation process. The anatomical connectivity patterns were obtained using dMRI tractography and were then used to perform the parcellation using a clustering algorithm. The functional connections, the cortical thickness gradient, and BigBrain data were used to verify the splitting trend. **(B)** Gene expression data acquisition and the process of differential expression analysis were performed using JuGEx, while the GSEA was employed to generate the ranked genes list and execute the gene enrichment analysis. **(C)** The process of studying how gene expression regulates both structural and functional connections during development using the PLS regression algorithm. The AHBA data was used to further explore the genetic factors of the division of the two subregions. **(D)** The functional connectivity patterns were obtained by calculating the correlation between brain regions using rfMRI data. The square shape represents the value of MCQ-K and the different colors represent different individuals. The relationship between the degree of similarity of the functional connectivity patterns between regions and the different behavioral expressions were explored using partial correlation. ^1^https://bigbrain.humanbrainproject.org

## Results

### Connectivity-based parcellation and validation

The optimal parcellation for each age bracket is shown in Figure 2A. The number of clusters was set from two to ten to obtain different parcellation patterns. The most appropriate number of partitions for each age bracket was assessed by multiple indexes, taking the similarity of both the topological and connectivity patterns into consideration. In terms of topological similarity, all of the indexes, Dice, NMI, and CV, achieved consistent results in that the local peak occurred when the number of clusters was six in the first three age brackets and seven in the adult one (Figure 2A for Dice and Figure S1 for NMI and CV). The SI index suggested that the local peak occurred at six clusters in the first age bracket. There was almost no local peak in the early-adolescence and post-adolescence stages, but seven clusters were identified in the adulthood stage. This suggested a clear transition, in that the optimal number of clusters changed from six to seven along with the age increase, as shown in Figure 2A. The IC index reflected the suitability of the group partition scheme across individuals. Similar to SI, the IC index in Figure 2A showed one of the local peaks at six clusters in the post-childhood stage and an approximate value of six and seven clusters for both the early-adolescence and post-adolescence stages. In young-adulthood, the minimum values of the IC were located at six clusters, which showed that a cluster number of six was definitely not a good option for the adulthood stage, although division seven performed only a slight better than division six according to this index. All of the indexes revealed an increasing trend in the optimal number of clusters with advancing age.

**Figure 2.**
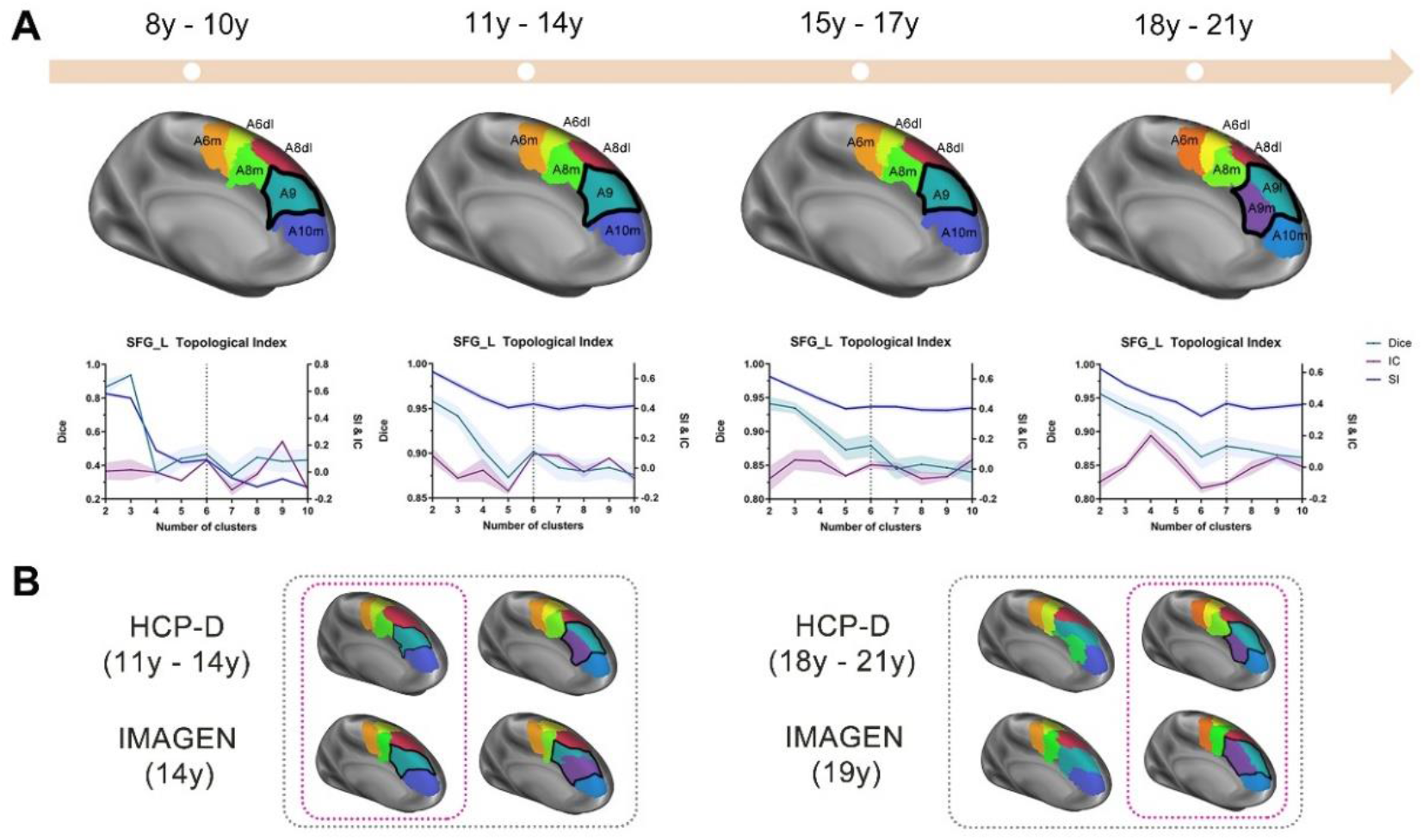
The parcellation results from childhood to adulthood and their validation from multimodal data. **(A)** The optimal partitioning pattern and corresponding indexes for each age bracket. From left to right, the age brackets are post-childhood, early-adolescence, post-adolescence, and young-adulthood. The indexes, Dice (cyan), IC (purple), and SI (blue), demonstrate the transformation of the parcellation pattern with age. Taken together, the six parcellation pattern was optimal for the first three age brackets but seven was optimal in adulthood. **(B)** Comparison between the six and seven parcellation patterns for different datasets. The most consistent parcellation patterns between the datasets are shown within the pink dashed lines for the adolescents (left) and adults (right).

The optimal parcellation pattern produced using another dataset, the IMAGEN dataset, is displayed in Figure 2B. The partitioning results clearly demonstrated that there was a more consistent parcellation pattern across different datasets when the number of partitions was six in the adolescents (14y). Conversely, the number was seven for young adulthood (19y) while the six-partition pattern showed a great deal of variation, especially in the A9 area. It seemed to stabilize as a seven-partition pattern after adulthood in a study by Fan et al. ^13^, in which they used data from subjects between the ages of 22 and 35 years old. The maximum probability maps of the two parcellation patterns at 14y and 19y are displayed in Figure S2 in the Supplementary material. There was no significant stability in the A9 region of the 14y 7-partition pattern, and the boundary within the A9 was different from that at 19y. By contrast, there were more stable subregions in the A9m and A9l at 19y, which indicates a higher level of partition consistency of the subregions in that group.

The overlap ratio (Table 1) of each subregion between the mean of the first three age brackets and adulthood was calculated to further study the relationship between the different subregions. We found that the overlap degree between the A9 regions in the childhood stage and the sum of the A9m and A9l regions in adulthood reached 0.82, suggesting that the A9m and A9l regions in adulthood were split in the A9 during development from post-childhood to adolescence.

**Table 1.** Dice of the subregions

|  |  |  |  |  |  |  |
| --- | --- | --- | --- | --- | --- | --- |
| Cluster number = 6 | A6m_L | A6dl_L | A8m_L | A9_L | A10m_L | A8dl_L |
| Cluster number = 7 | A6m_L | A6dl_L | A8m_L | A9m_L+ A9l_L | A10m_L | A8dl_L |
| Dice | 0.80 | 0.85 | 0.86 | 0.33+0.49 | 0.75 | 0.61 |

The data from different modalities were used to further verify the trend in the division. The resting-state functional connectivity boundary maps are shown in Figure S3A, in which the ages are arranged from younger to older. The first two maps show no clear dividing line at the boundary between the A9m and the A9l in adulthood, whereas the last two maps showed more overlap of the boundary obtained by the boundary mapping method (using fMRI data), and the clustering method (using dMRI data) had a greater degree of coincidence, which suggests that the differences between the two regions became more pronounced.

Because there was no suitable method for selecting the coordinate positions of the specific points, five points along the vertical, perpendicular to the boundary between the A9l and the A9m, were picked manually. Then the point-taking process was repeated five times along the rostral to the caudal direction in the SFG, and an averaging operation was performed on the five collections. Finally, we gathered the values from the points perpendicular to the dividing line, that is, the gradient of the thickness (Figure S3B). The gradient values of points near the boundary increased with increasing age. In other words, the regional differences between each side at the boundary became more significant with age, which supports our conjecture about the division of subregions in the A9l and A9m. The distribution of gray matter density values in the A9l and A9m areas is shown in Figure S3C. Significant differences between distributions suggest that the cell types differed between the A9l and the A9m.

### Different developmental trajectory of connectivity patterns in the medial-lateral dorsal PFC

To reveal the reasons for the different cluster numbers in the A9l and A9m between the adolescents and the young adults, we explored the connectivity patterns between 246 target subregions and the two regions separately for each age bracket (Figure S4 and Figure S5). The results showed increasing differences in the connectivity patterns between the two subregions with age (Figure 3A). The differences in the connectivity patterns between the A9m and A9l regions were mainly in their connection with the cingulate gyrus, most of the subregions of the frontal lobes, and some of the regions of the hippocampus, basal ganglia, and thalamus in the subcortical nuclei in the left hemisphere. In the right hemisphere, the different regions were located in areas of the frontal lobe and limbic lobe. What is noteworthy is that the connections between the source regions (A9m and A9l) and some of the target regions (the parietal lobe of the left hemisphere and the frontal lobe, parietal lobe, and occipital lobe of the right hemisphere) became more diverse with age, which suggests that the splitting of the A9 may be caused by changes in the functional coupling of the A9m and A9l with these regions.

**Figure 3.**
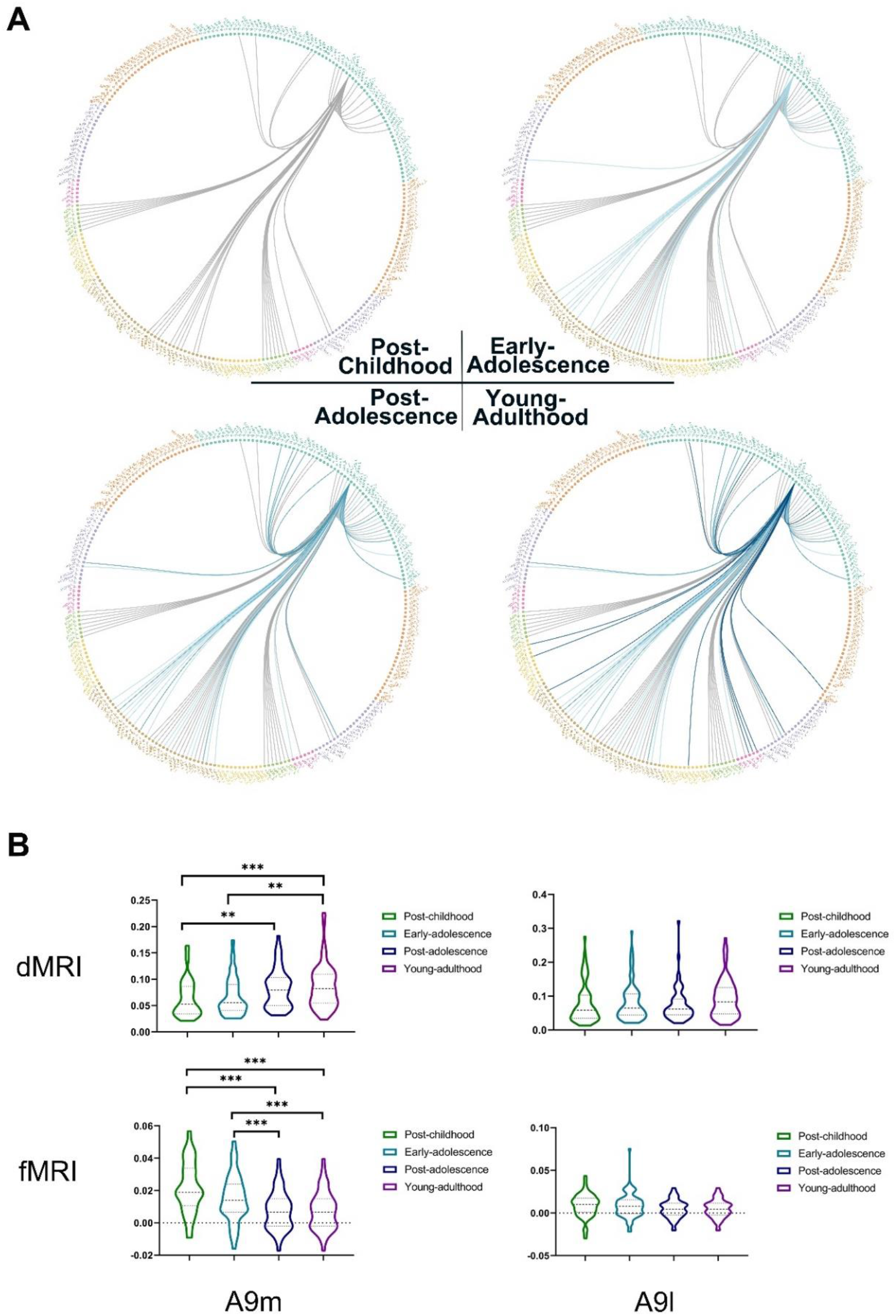
Changing connections with age. (A) Differences in the connections between the A9m and A9l areas in each age bracket. (B) Results from statistical analyses of the dMRI and fMRI data.

The results of the ANOVA analyses on the A9m and A9l (with the parcellation of the Brainnetome atlas as target areas) are shown in Figure 3B. Significant structural connection differences in the ANOVA analyses between different age groups were demonstrated in the A9m (*F* = 18.01; *p* < .01; *d* = 3). Considered separately, significant differences occurred between the post-childhood and post-adolescence groups (*p* < .01, FWE-corrected), the post-childhood and young adulthood groups (*p* < .01, FWE-corrected), and the early-adolescence and post-adolescence groups (*p* < .01, FWE-corrected). No significant results were observed in the A9l, which suggests that more changes had taken place in the A9m than in the A9l during the developmental process. Similar results were found from the fMRI data. Significant functional connection differences between different age groups in the ANOVA analyses were also demonstrated in the A9m (*F* = 14.65; *p* < .01; *d* = 3). Significant differences also appeared between the post-childhood and post-adolescence groups (*p* < .01, FWE-corrected), post-childhood and young-adulthood groups (*p* < .01, FWE-corrected), early-adolescence and post-adolescence groups (*p* < .01, FWE-corrected), and early-adolescence and young-adulthood groups (*p* < .01, FWE-corrected). In addition, ANOVA analyses were performed on the fMRI data using the 7 networks defined by Yeo et al. ^51^ as the targets. Significant connection differences between the ages occurred between A9m and the limbic, fronto-parietal, and default networks (Figure S6). Only the connection between the A9l and the ventral attention network were found to be significant between ages (Figure S7).

### Association between genetic variation and the segmentation of the medial-lateral PFC

A differential expression analysis of the genes revealed differential SYT10 gene expression between the A9m and the A9l. SYT10 is a protein-coding gene related to protein-protein interactions at synapses and transmission across chemical synapses that play a role in exocytosis and the regulation of dopamine secretion. The gene expression level of SYT10 in the dorsal frontal cortex (DFC) and medial frontal cortex (MFC) changes from early childhood with the difference peaking during adolescence, suggesting that increased dopamine secretion and protein activity in the MFC during development may lead to more changes in the medial prefrontal lobe than in the lateral in the maturation process ^68^.

The gene enrichment analysis showed more intense enrichment in the A9m area than in the A9l for the regulatory target gene sets, which represent potential targets of regulation by transcription factors or microRNAs. Oncogenic signature gene sets, which represent signatures of cellular pathways that are often dis-regulated in cancer ^57^, were also found to be enriched in the A9m but not in the A9l area, implying that abnormal gene expression in the A9m may contribute to abnormal cell division and proliferation, which may, in turn, cause many developmental diseases.

We further explored the relationship between the gene expression and the age-related connectivity pattern for each subregion (A9m and A9l) in the dMRI and fMRI data using PLS (Figure 5A, 5B, respectively). Only the valid components that captured the greatest variance in the age-related structural and functional connections, defined as s-PLS1 and f-PLS1, respectively, are shown in Figure 4. We focused on the detailed analysis of these components here in the main text. In the A9m, the s-PLS1 captured 13.5% of the age-related structural connectivity (SC) (*p* < .05), while the s-PLS1 explained 15.5% (*p* < .05) in the A9l. The f-PLS1 in the A9m and A9l respectively explained 17.4% and 17.5% of the age-related FC variance (both *p* < .001). We ranked the normalized weights of these validated components based on univariate one-sample *Z* tests to find the overexpressed (*Z* > 5) and underexpressed (*Z* < −5) genes lists. Then, 13 overexpressed genes and 8 underexpressed genes were identified as being related to the development of age-related SC in the A9m, and 31 overexpressed genes and 17 underexpressed genes were identified in the A9l (all *p*-FDR < .005; Figure S8 A and Figure S8 B). In the A9l area, 39 genes and 53 genes that were related to the development of functional connections were respectively found to be overexpressed and underexpressed (Figure S8 C). Therefore, there were too few candidate genes to be suitable for an enrichment analysis. However, for the f_PLS1, 131 overexpressed genes and 313 underexpressed genes were found to be significantly associated with age-related FC in the A9m (Figure 5C), which was significantly enriched for the underlying GO annotations. What is noteworthy is that genes for the positive regulation of phospholipid biosynthesis were found to be the ones that were enriched relative to developing connections in the A9m, which suggests that specific gene expression patterns influence regional parcellation patterns by influencing neurophysiological processes, such as myelination (Figure 5D).

**Figure 4.**
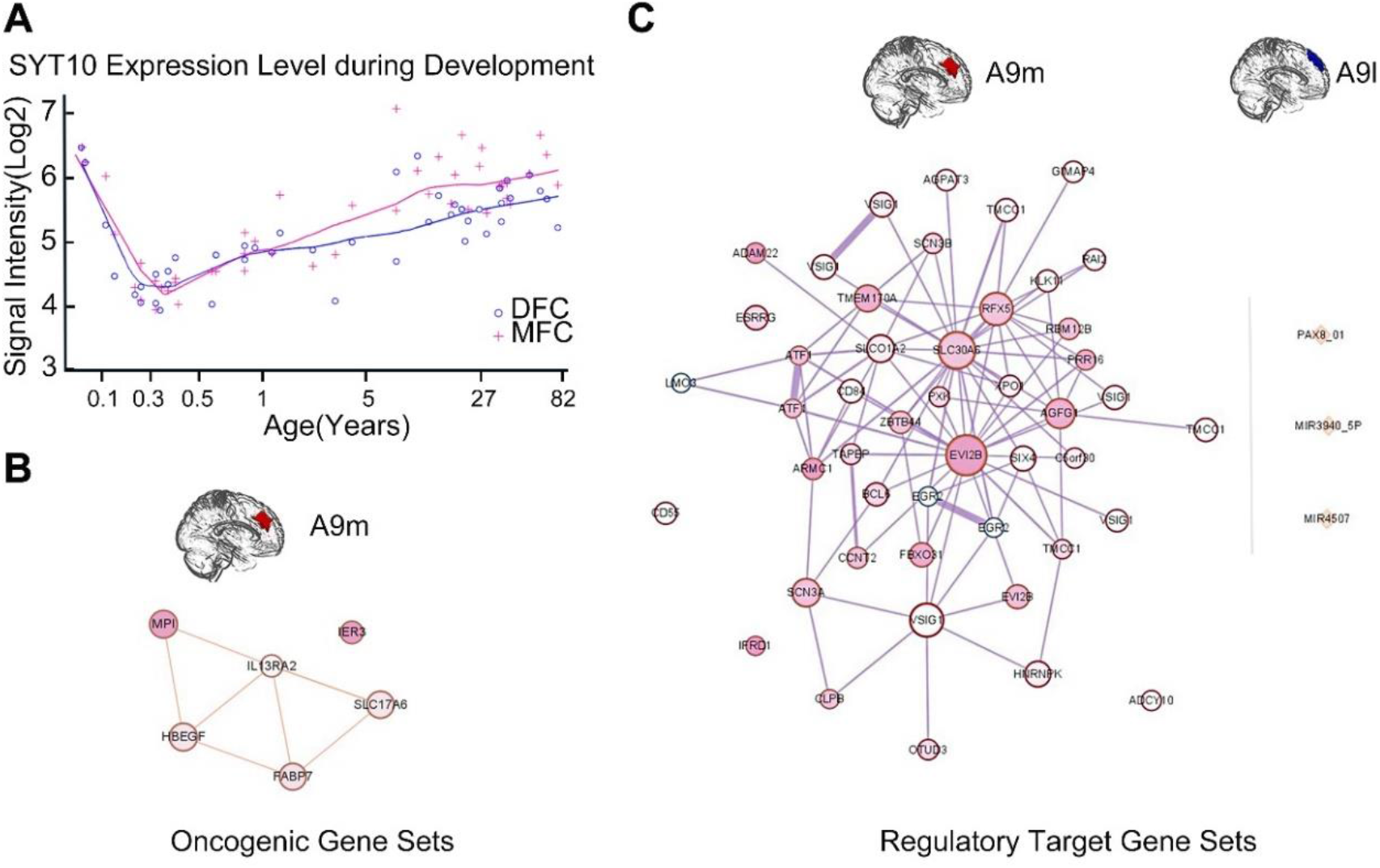
Results of differential expression analysis and gene enrichment analysis. **(A)** The pink and blue lines show the changes in gene expression during development in the MFC and DFC, respectively. **(B)** The thickness of the line and the size and color of the dots represent the strength of the connections between the gene sets, the number of genes in each gene set, and the significance of the p value (red for up-regulated genes and blue for down-regulated genes), respectively.

**Figure 5.**
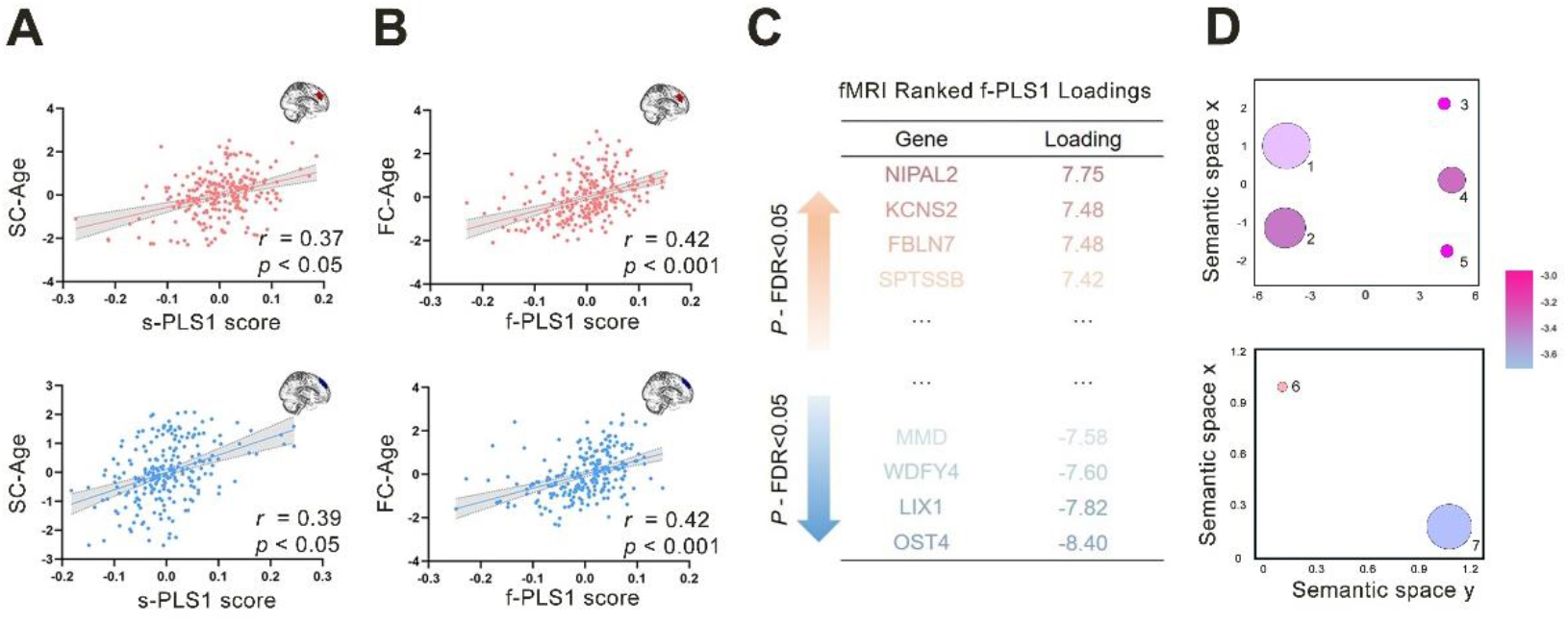
Coupling between age-related connections and gene expression. The scatterplot of the PLS valid components scores and both **(A)** structural and **(B)** functional connections. The correlation of the A9m region is in pink, while the blue color was used for the A9l. SC, structural connection; FC, functional connection. **(C)** Ranked valid components loadings of the A9m. **(D)** Physiological processes that showed significant enrichment. 1, regulation of gene expression; 2, regulation of transcription and DNA-templated synthesis; 3, negative regulation of cell communication; 4, negative regulation of response to stimulus; 5, negative regulation of signaling; 6, positive regulation of phospholipid biosynthetic process; 7, positive regulation of chondrocyte differentiation. Bubble color indicates the p value and size indicates the frequency of the GO term in the underlying GOA database ^69^.

### Maturation correlation between behavior and functional connectivity of the A9m

A behavioral relevance study was carried out using the IMAGEN dataset. A meta-analysis showed that both the A9m and the A9l were involved in the inference process. The difference was that the A9m was more related to preference and strategy, while the A9l had more to do with socialization and cognition. Based on prior knowledge, the collected data from the MCQ were further processed to reveal the relationship between the A9m and A9l subregions and changes of behavior during development. In the MCQ, a higher K value indicates a stronger impulsivity preference for immediate but lower value rewards. The result showed that the greater the similarity between a subject’s functional similarity at A9m and A9l the smaller the K values in the same individual at 19 years old. (Figure 6B), that is, subjects that were highly similar in the two subregions at 14 had a higher capacity for non-impulsivity in the young-adulthood stage.

**Figure 6.**
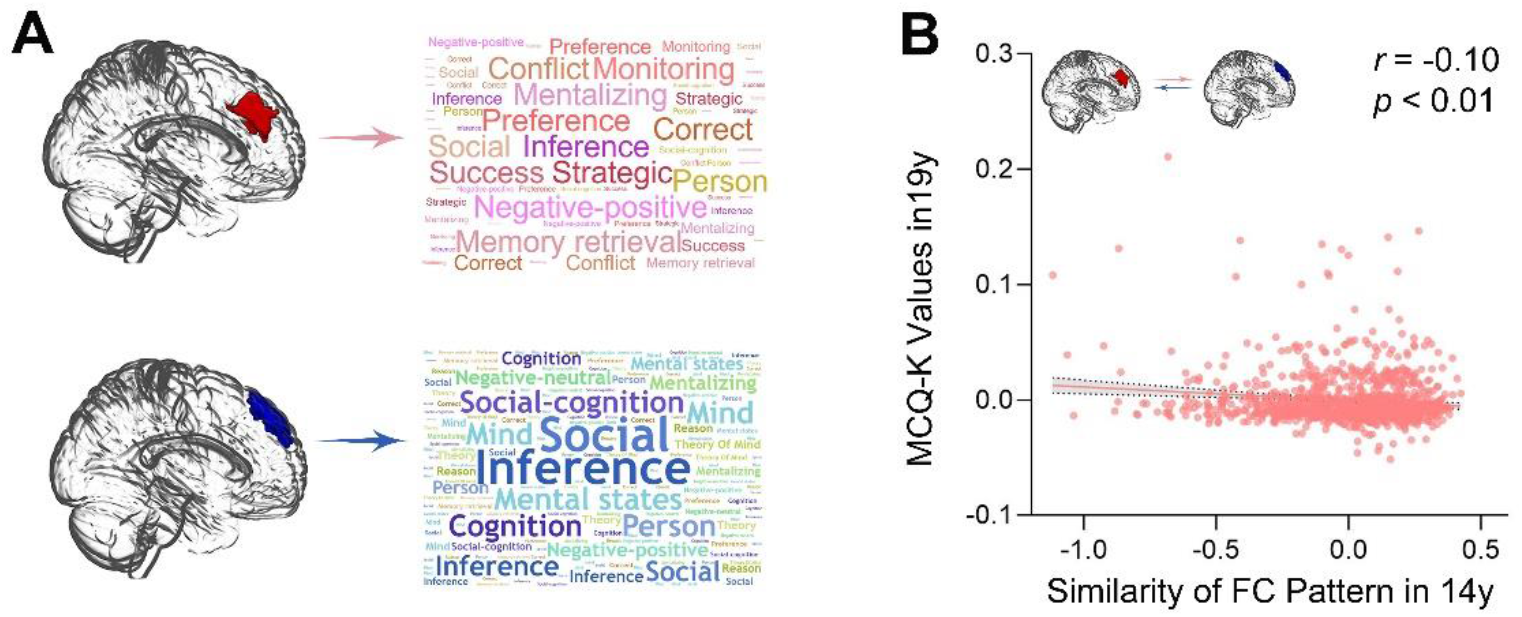
Connections change with age. **(A)** Results of meta-analysis of the A9m and A9l. **(B)** Correlation between the similarity of the functional connection and the behavioral value assessed by the MCQ-K.

## Discussion

Childhood and adolescence are two periods with extensive cortical maturation in the PFC, causing cortical patterning to be substantially dynamic across the developmental stages and shaped by the interplay of intrinsic and extrinsic mechanisms. In the present study, we delineated the human cortical parcellation trajectory of the SFG based on anatomical connectivity profiles. Specifically, we observed that the A9 region separated into medial and lateral subregions with the advent of adulthood, as validated by several cohorts with multi-modal data that included gradients in cortical thickness, functional connectivity patterns, cytoarchitectonic maps, and transcriptional profiles. This separation revealed by connectivity contrasts may occur gradually with age, reaching a specific boundary at the transition from adolescence to young adulthood. Accordingly, we found that the differential gene expression patterns between the areas that showed a tendency to segregate originated in toddlers and widened increasingly distinctly during adolescence. These changes may be involved in the process of medial-lateral differentiation in the dorsal PFC. The gene-driven transformations of connectivity patterns during maturation were also engaged in the arealization process, especially as indicated by more varied connections with the thalamus. In addition, the transformation of connectivity patterns appears to have an effect on development and behavior that is mostly concerned with a young person’s persistence and the degree to which they show non-impulsiveness.

### Developmental trajectory of arealization and connectivity patterns

Unquestionably, brain characteristics change dynamically during development. In the context of a brain atlas, this is expressed as differential arealization patterns. In the dorsal prefrontal cortex, significant arealization differences were found between minors and adults, demonstrated as the segregation of the A9 area into A9m and A9l sub-areas, suggesting the importance of constructing specific brain maps for children. Changes in neuroanatomy potentially contribute to this differentiation. Our findings also demonstrated that changes in cortical thickness gradients accompany changes in arealization patterns. Previous studies also found that during development the prefrontal cortex undergoes a remarkable neuroanatomical transformation, including a reduction in synaptic and neuronal density, a growth of dendrites, and an increase in white matter volume 10. In addition, this separation arose from increasing differences in the connectivities of the two subregions with age and may begin as early as post-adolescence or even earlier in early-adolescence but does not show up until young adulthood in the form of a parcellation pattern. Interestingly, previous research indicated that the medial PFC actively enters a prolonged myelination cycle at the time of adolescence, during which it increases in plasticity and, thus, reflects adaptations to the ever-changing environment ^70^. By impacting the structural connections, this myelination cycle may be the essential trigger for the developing arealization pattern.

In detail, the change in connectivity pattern manifests as a differentiation in the target areas arrived at by the white matter fiber bundles. The differences between the connected target subregions of the A9m and A9l are generally in the frontal lobe, cingulate gyrus, and subcutaneous nuclei. Specifically, the increasing difference in connections to the thalamus with age was especially obvious and hinted that area-specific maturation-related signal inputs from extrinsic areas, particularly the thalamus, may play an important role in the final maturation regionalization pattern. Consistent with our finding, previous studies also showed that the establishment of boundaries between cortical areas depended on thalamocortical input ^71,72^. In addition, the increasingly different connections to the insula with the advent of adulthood may also contribute to the formation of the final boundary. The variation in the target region reached by the seed regions also suggests a shift in the function of the developing seed regions. Given that intricate activities depend on the collaboration between different areas and that brain areas generally correspond to dissimilar functions, the age-dependent discrepant connectivity patterns of the two areas suggest their gradual functional differentiation, especially in decision-making, inhibition ability, and emotional and cognitive development ^73–76^. These functions correspond to target areas in the cingulate cortex, basal ganglia, hippocampus, thalamus, and regions that collaborate with them. In general, this separation suggests that the functions of the dorsal prefrontal cortex shift from a relatively general ability to more differentiated cognitive abilities ^77^, a process which is likely constrained by changes in the underlying connectivity architectures during development.

The later or earlier maturation of regional separation patterns also correlates with individual behavior patterns. Compared with younger children, older adolescents have different brain activity patterns related to intertemporal decision-making when choosing between immediate and delayed rewards ^78^. In this study, our results suggest that individuals who develop late tend to have a higher ability to achieve delayed gratification, which means that earlier maturation of the brain is not necessarily a good thing for teenagers. Related research showing that earlier maturation of brain features may increase the likelihood of engaging in substance abuse behaviors also provides evidence for this view ^79^. Delayed discounting has been studied extensively as a key personality trait that is important in enabling adolescents to transition from children to adults. Previous study has shown an age-related preference for immediate versus delayed rewards, which was associated with white matter integrity in the left frontal lobe ^80^. Regional homogeneity patterns of the dmPFC were also found to contribute to the delayed discounting rate ^81^. However, we have offered a potential new perspective for exploring the relationship between changes in connectivity patterns between the dlPFC and dmPFC during development and the delayed discounting rate. What is more, adolescents with abnormal MCQ scores may be at risk for mental disorders and have excessive sensitivity to rewards. Many patients with drug and alcohol dependence cannot tolerate the temptation of smaller immediate rewards rather than waiting to get larger rewards at a later date ^82^. Thus, this study of brain arealization in each age stage may contribute to the early detection of mental diseases and the prevention of drug addiction.

### Validation of medial-lateral separation using multi-index, cross-modality, cross-sectional, and longitudinal data cohorts

One challenge when parcellating using multimodal imaging data is how to determine the proper number of partitions. One strategy is to integrate multimodal information as a unified feature before partitioning ^49^. This method has the advantage of avoiding a non-uniform number of partitions from multi-model data. However, this same strength limits the interpretability and relevance of such a map, as the ensuing parcellation might be biased by a priori knowledge and expectations about brain organization ^11^. Here, we used separate indexes that can represent the overlap of voxel within parcels, cluster consistency, and the partition of individual variability to measure the appropriateness of the partition model. The most appropriate parcellation pattern was defined as the one that performed well on the most indexes. This was the 6-partition pattern in childhood and the 7-partition pattern in adulthood.

For the cross-modalities verification, after determining the parcellation pattern mainly based on anatomical white matter pathways, we further tested whether this pattern could be supported by the following features: functional connections based on fMRI, cortical thickness, and cytoarchitectonic and genetic information. Limited by the data provided by each mode and considering that each neurobiological characteristic represents a unique view of brain tissue, the method and result of verification for each mode was not the same. Using the boundary mapping method for the fMRI data, we found that the boundary between the A9l and A9m became more significant with age; using cortical thickness, we found a differential gray matter density distribution between the medial and lateral subregions; using gene enrichment analysis, the two areas also showed heterogeneous gene expression patterns in adulthood, which indicated a physiological functional heterogeneity between the two areas. Thus, each type of multimodal data provided evidence for segregation in the dorsal prefrontal cortex.

To verify the parcellation results using multiple cohorts, we used another dataset (the IMAGEN dataset) to identify the parcellation pattern. The optimal number of partitions for different age groups, that is, the 6-partition pattern in childhood and the 7-partition pattern in adulthood, had the best overlap between the different data sets. Although the parcellation patterns of the two datasets were not the same, this might be due to the limited resolution and b-value of the dMRI data parcellation of the IMAGEN dataset. Nevertheless, the similar segregation-tendency with age in the A9 area in the IMAGEN dataset is noteworthy and further validated our findings.

In particular, when we performed separate gene enrichment analyses in the A9m and A9l regions, we found that many of the gene sets that were concentrated only in the A9m region were involved in myelination. For instance, among the enriched oncogenic signature gene sets, HBEGF was found to be widely distributed in the cerebral neurons and neuroglia ^83^. Also, FABP7 plays an important role in the establishment of radial glial fibers during brain development ^84^ and regulates the proliferation and differentiation of oligodendrocyte lineage cells ^85^. The enriched gene in the regulation of target genes, FBXO31, was also found to take part in dendrite growth and neuronal migration ^86^. Thus, the oligodendrocyte lineage cells participate in forming myelin sheaths and supporting axons metabolically ^87^, whereas some other physiological process may indirectly influence the formation of developmental regionalization.

In addition, using differential gene analysis between A9m and A9l, only SYT10 was observed to be significantly differentiated, and its expression level was much higher in the A9m. The GO-Molecular function of SYT10 is related to calcium-dependent phospholipid binding ^88^. When associating gene expression with age-related connections using PLS, the genes included in the f-PLS1 were also found to be enriched in the process of phospholipid biosynthesis, which is also associated with neural growth. Interestingly, given that SYT10 is one of the most enriched genes in the MFC during adolescence ^89^, different expression trajectories of the SYT10 were found between the medial and lateral areas of the PFC. In detail, such a difference begins at toddlerhood and is increasingly distinct throughout successive developmental time points with the peak discrepancy around early adulthood. Convergent evidence from animal ^90^ and human studies ^91 92^ has demonstrated a genetic influence on the specification of distinct cortical regions. Our findings echoed these studies, and together they suggest that such region-specific transcriptional patterns may underlie long-lasting physiological processes that contribute to the construction and functional specializations of brain areas.

### Methodological considerations

Nonetheless, more work is needed to gain a better understanding of the developing brain. Changes in regionalization patterns in other subregions still remain obscure, perhaps because the target areas of the ROI-based connections were often based on adult atlases in most of the present developmental studies, but such studies are not entirely suitable for children’s studies. We plan to complete the parcellation of the remaining brain regions in the near future. Additionally, due to a lack of longitudinal data over a wider age range, the developmental characteristics of neither the earlier nor later stages of life are still unclear. The ongoing ABCD project is collecting longitudinal data for at least 10 years starting at ages 9 to10 ^93^. This data will allow us to implement more complete research in the future. With respect to cytoarchitectonics, no study involving a cytoarchitectural atlas of the whole brain throughout the entire development process has been completed. The same drawback exists with respect to a gene expression atlas; the Julich-Brain atlases are still partially incomplete ^94^. In the near future, we plan to delve deeper into the mechanisms by which genes regulate brain activity and study this using monozygotic and dizygotic twins utilizing bivariate twin models.

### Conclusions

In summary, our research provides new insight into the cortical regionalization patterns of the developmental human SFG, suggesting that a functional separation of the dmPFC and dlPFC exists. By exploring the developmental trajectory of the connectivity patterns in the seed areas, as well as by a genetic analysis, we were able to identify the separation of the dmPFC and dlPFC and found a potential mechanism causing this separation. The cortical regionalization patterns established during early development seem to be affected by gene expression patterns and refined by behavior-related transformations of the connectivity patterns during later development to generate the mature cortical areas. These results provide new evidence for the developmental changes in the prefrontal lobes and shed new light on their impact on individual development.

## Supporting information

supplemental tables and supplemental pictures

## Data availability

Data from both the Lifespan Human Connectome Project in Development and the Adolescent Brain Cognitive Development project can be requested from the NIMH Data Archive (NDA). The IMAGEN data can be found at https://imagen2.cea.fr/database/. The human gene expression data are available in the Allen Brain Atlas (https://human.brain-map.org/static/download).

## Code availability

The HCP-Pipeline can be found at https://github.com/Washington-University/HCPpipelines. The neuroimaging preprocessing software used for the other datasets is freely available (FreeSurfer v6.0, http://surfer.nmr.mgh.harvard.edu/, and FSL v6.0, https://fsl.fmrib.ox.ac.uk/fsl/fslwiki). The gene expression code analysis can be found at https://figshare.com/articles/code/TBTA_zip/11101472. Gene enrichments were analyzed using GORILLA, which can be found at http://cbl-gorilla.cs.technion.ac.il/ and were followed by visualization using REVIGO at http://revigo.irb.hr/. The brain maps were presented using Surf Ice (v2.0, https://www.nitrc.org/projects/surfice/) and WorkBench (v1.5.0, https://www.humanconnectome.org/software/get-connectome-workbench).

## Acknowledgments

We are very grateful to all the participants who contributed to this article. R. E. Perozzi and E. F. Perozzi provided assistance with English language and editing.

## Author contributions

T.J. led the project and the concept design. W.L. carried out the data processing. W.L. and T.J. analyzed the data, created the figures and wrote the manuscript. T.J., L.F., W.S., H.W., J.L., Y.C., K.L., L.C. and Y.L. provided crucial advice for the study. All of the co-authors participated in discussions of the results and revision of the manuscript.

