## supplemental tables and supplemental pictures for "Anatomical connectivity profile development constrains medial-lateral topography in the dorsal prefrontal cortex"

### Methods

#### 1.1 Definition of developmental age brackets

Based on the way that some influential organization, such as Healthy Children, World Health Organization, American Academy of Pediatrics, and American Academy of Child and Adolescent Psychiatry, supplemented by articles related to development (Williams, Ponesse, Schachar, Logan, & Tannock, 1999; Xia et al., 2018), defined age, we divided all ages of the HCP-D data (8-21y) into 4 stages, post-childhood (8-10y), early-adolescence (11-14y), post-adolescence (15-17y) and young-adulthood (18-21y) (Table S1). Because of individual differences, different but similar age group divisions had little influence on the group parcellation patterns.

**Table S1.**

| Organizations [a] or references [b] | Post-Childhood | Early-Adolescence | Post-Adolescence | Young-Adulthood |
|---|---|---|---|---|
| Healthy Children [a] | 8-9 | 10-13 | 14-17 | 18-21 |
| American Academy of Pediatrics [a] | 8-10 | 11-14 | 15-17 | 18-21 |
| Association of Maternal & Child Health Programs [a] | 8-9 | 10-14 | 15-17 | 18-21 |
| American Academy of Child and Adolescent Psychiatry [a] | 8-10 | 11-13 | 14-18 | 19-21 |
| World Health Organization [a] | 8-9 | 10-15 | 14-17 | 16-21 |
| Williams, Benjamin R., et al, 1999 [b] | 8 | 9-12 | 13-17 | 18-21 |
| Xia, Cedric Huchuan, et al, 2018 [b] | 8-10 | 11-13 | 14-16 | 17-19 |
| Consensus age range divisions | 8-10 | 11-14 | 15-17 | 18-21 |

All divisions are made on a yearly basis. [a] Organizations, [b] References.

#### 1.2 Discarded ROIs from the Brainnetome Atlas that had no samples in the Allen Brain Atlas

**Table S2.**

| | ROIs |
|---|---|
| Right hemisphere | *A6dl, dorsolateral area 6*<br>*A8vl, ventrolateral area 8*<br>*A44d, dorsal area 44*<br>*A45c, caudal area 45*<br>*A44op, opercular area 44*<br>*A6cvl, caudal ventrolateral area 6*<br>*A1/2/3ll, area1/2/3 (lower limb region)*<br>*TI, area TI (temporal agranular insular cortex)*<br>*A7pc, postcentral area 7*<br>*OPC, occipital polar cortex*<br>*msOccG, medial superior occipital gyrus* |

### Results

#### 2.1 Verifying parcellation reliability

As shown in Figure S1, both the NMI and CV indexes demonstrated that the 6-partition pattern was the most suitable for the early-adolescence and post-adolescence groups and that a 7-partition pattern was most suitable for the young-adulthood group. Both indexes seem to indicate that the 3, 6, and 8 partion patterns were suitable for the post-childhood group; however, after considering other indexes, we finally chose the 6-partition pattern for the post-childhood group.

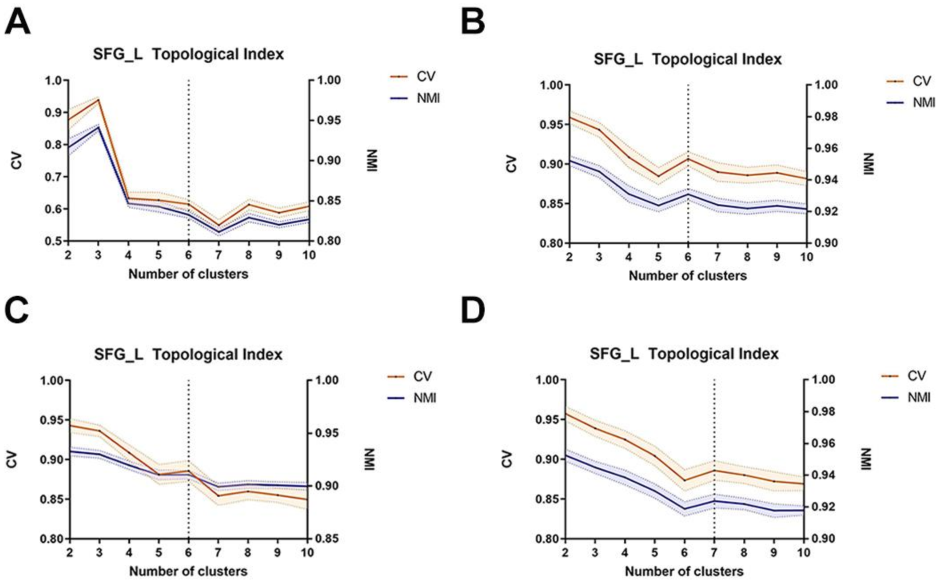

***Fig. S1 Verification of topological consistency using NMI and Cramer's V (CV). A, B, C, D** show the topological indexes for the post-childhood (8-10y), early-adolescence (11-14y), post-adolescence (15-17y), and young-adulthood (18-21y) groups, respectively.*

The maximum probability maps below display the consistency of the partition patterns between groups at ages 14y and 19y, in which the sharpness of the color values reveals the consistency within groups. The red color represents stable subregions of the partition across subjects, the green color represents the boundary, and the blue color is the unstable area in between. The higher regional consistency in the A9 subregion at age 14 suggests the rationality of the 6-partition pattern at this time; in contrast, the 7-partition pattern revealed almost no stable region in Figure S2 in the A9. At age 19, the 6-partition pattern was not a good choice due to significant inconsistencies between the data sets, but the 7-partition pattern was more acceptable.

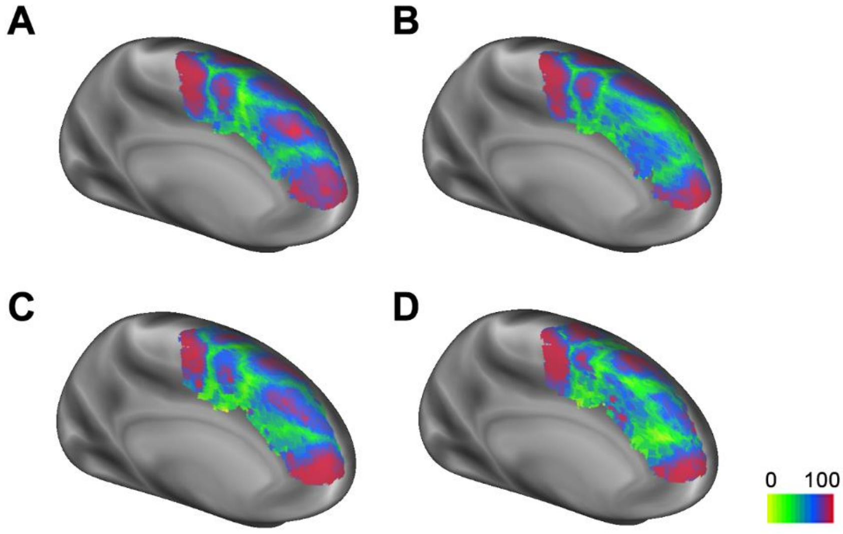


***Fig. S2 The maximum probability maps. (A)** The 6-partition pattern at 14y. **(B)** The 7-partition pattern at 14y. **(C)** The 6-partition pattern at 19y. **(D)** The 7-partitioning pattern at 19y.*

The gradient-based parcellation based on fMRI used the watershed-by-flooding algorithm. We referred to the process by which other researchers had established a whole brain parcellation pattern (Cohen et al., 2008; Evan et al., 2016; Wig, Laumann, & Petersen, 2014). In each cortical hemisphere, the gradients were averaged and smoothed across subjects on the brain surface, and watershed-based edge detection was applied to generate a group-average edge map for each vertex. The group-average edge density map was generated by averaging the edge maps.

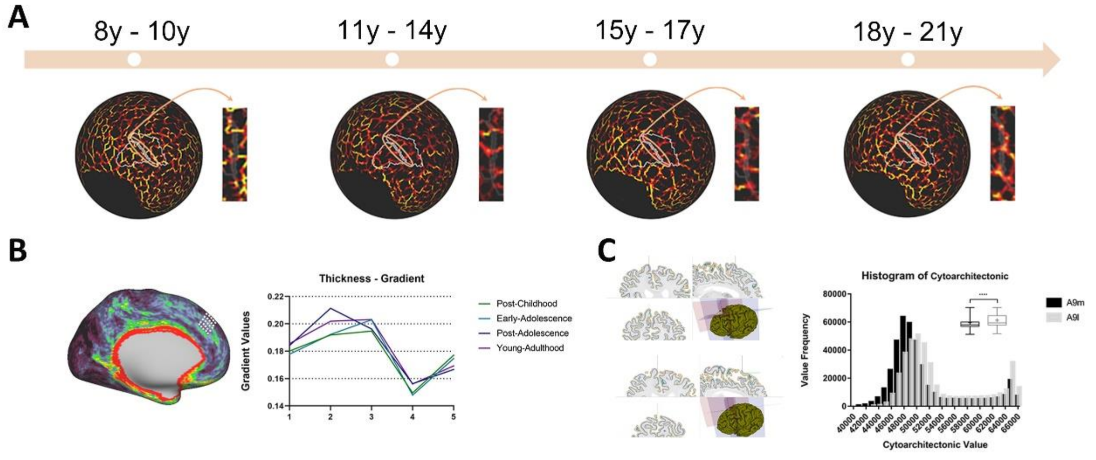


***Fig. S3 Parcellation results from childhood to adulthood and their validation from multimodal data. (A)*** *The boundary mapping produce using fMRI data. The area where the subdividing lines are located is shown separately.* ***(B)*** *The gradient changes in cortical thickness increased with age.* ***(C)*** *Comparison of the cytoarchitectonic values between the A9m and A9l subregions in adulthood.*

#### 2.2 Changes in connection patterns with age

The connectivity pattern of the two subregions by age is shown below. The connectograms were obtained under the same processing flow and parameters.

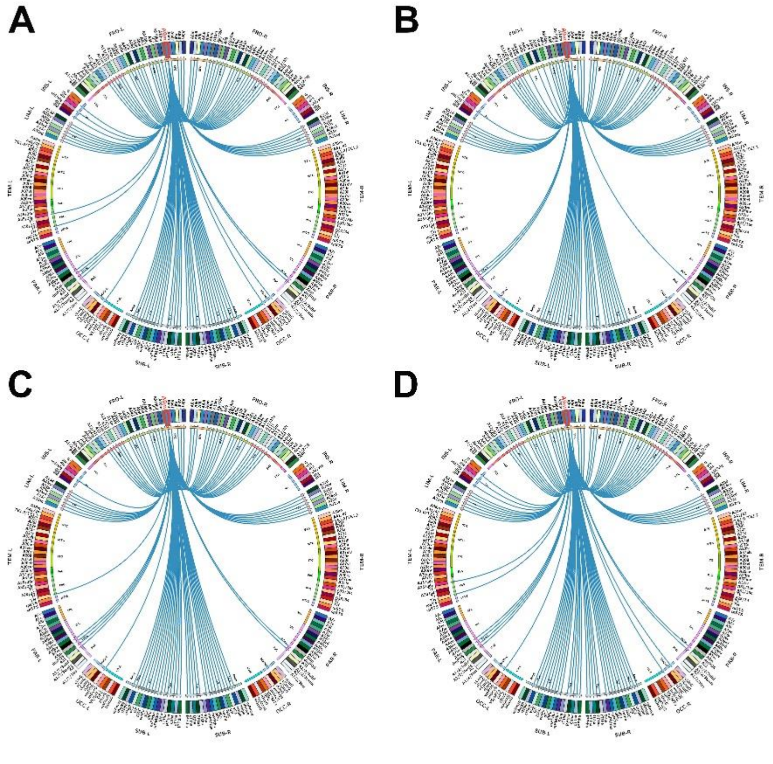

***Fig. S4 Connectograms of the A9m. A, B, C, D** show the connectograms of the A9m for the post-childhood (8-10y), early-adolescence (11-14y), post-adolescence (15-17y), and young-adulthood (18-21y) groups, respectively.*

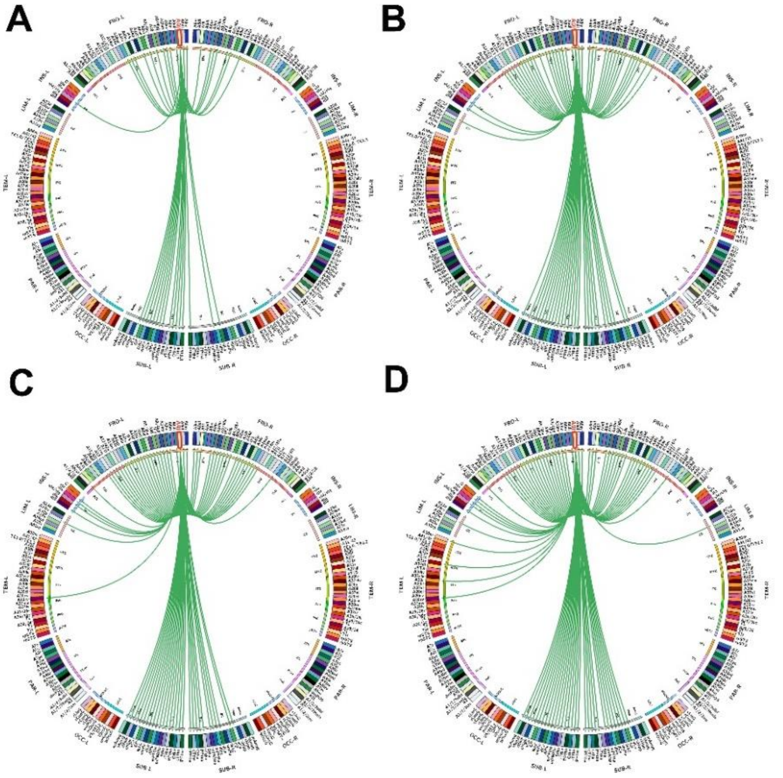


***Fig. S5 Connectograms of the A9l. A, B, C, D** show the connectograms of the A9l for the post-childhood (8-10y), early-adolescence (11-14y), post-adolescence (15-17y), and young-adulthood (18-21y )groups, respectively.*

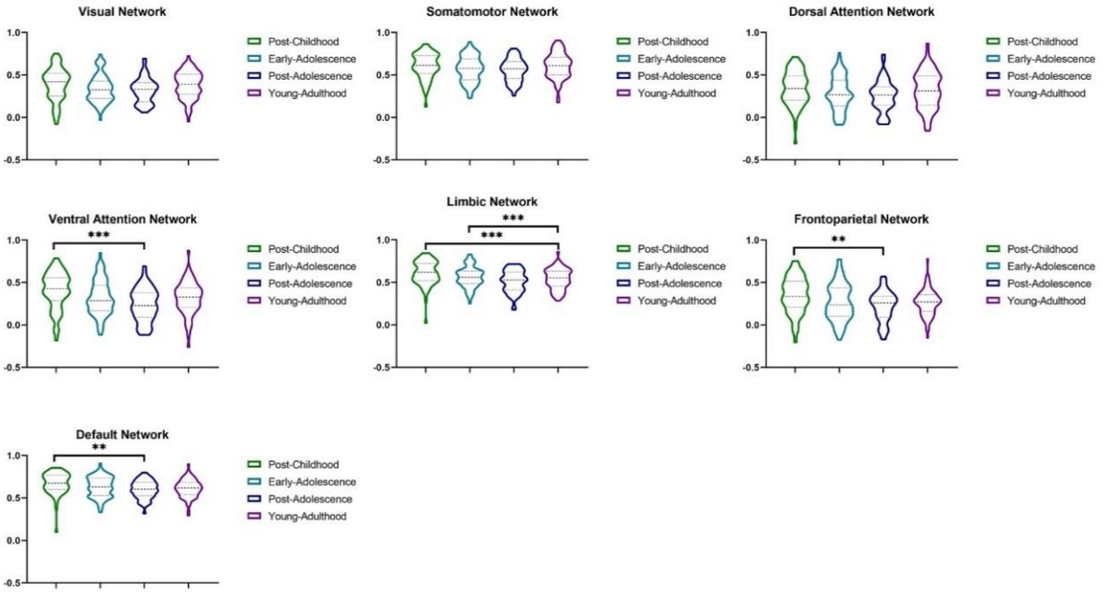

***Fig. S6 ANOVA analyses between different age stages for the functional connections of the A9m.*** *The target areas are 7 networks from Yeo. Only significance with p < .01 is shown.*

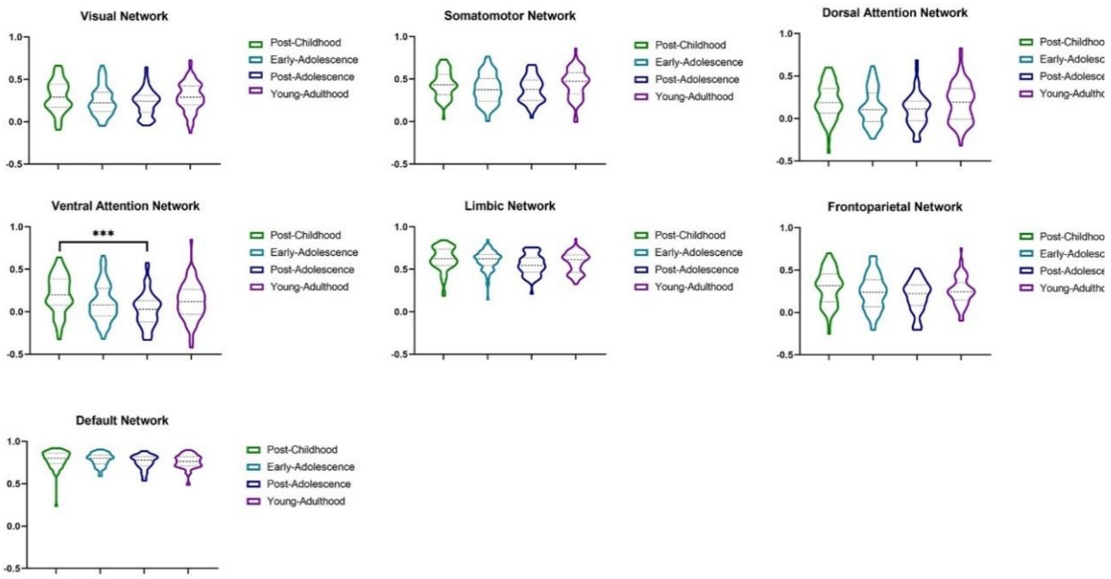


***Fig. S7 ANOVA analyses between different age stages for the functional connections of the A9l.*** *The target areas are 7 networks from Yeo. Only significance with p < .01 is shown.*

#### 2.3 Ranked PLS1 loadings of dMRI and fMRI

**A** DTI Ranked PLS1 loadings

| | Gene | Loading |
|---|---|---|
| P - FDR<0.05 ↑ | GRIK1 | 5.80 |
| | CHRNA2 | 5.68 |
| | MANBA | 5.47 |
| | RGS16 | 5.40 |
| | … | … |
| | … | … |
| P - FDR<0.05 ↓ | CHP2 | -5.32 |
| | NUDT1 | -5.35 |
| | EEF1AKMT4 | -5.67 |
| | GRID1 | -5.83 |

**B** DTI Ranked PLS1 loadings

| | Gene | Loading |
|---|---|---|
| P - FDR<0.05 ↑ | CDADC1 | 6.77 |
| | PRPSAP2 | 5.95 |
| | NOV | 5.87 |
| | KCNK10 | 5.52 |
| | … | … |
| | … | … |
| P - FDR<0.05 ↓ | TRMT6 | -5.76 |
| | AMY2A | -5.93 |
| | SH2B1 | -6.04 |
| | TSC2 | -6.22 |

**C** fMRI Ranked PLS1 loadings

| | Gene | Loading |
|---|---|---|
| P - FDR<0.05 ↑ | SOWAHB | 6.25 |
| | FBLN7 | 6.16 |
| | STX19 | 5.97 |
| | PP12613 | 5.96 |
| | … | … |
| | … | … |
| P - FDR<0.05 ↓ | ACSBG1 | -6.29 |
| | CYBRD1 | -6.32 |
| | SOX9-AS1 | -6.54 |
| | SELENBP1 | -6.71 |

***Fig. S8 Ranked valid components loadings. (A)*** *The overexpressed and under-expressed genes found to be related to the development of structural connections in the A9m.* ***(B)*** *The overexpressed and under-expressed genes found to be related to the development of structural connections in the A9l based on DTI.* ***(C)*** *The overexpressed and under-expressed genes found to be related to the development of functional connections in the A9l based on fMRI.*
